# Cullin-5 adaptor SPSB1 controls NF-κB activation downstream of multiple signalling pathways

**DOI:** 10.1101/634881

**Authors:** Iliana Georgana, Carlos Maluquer de Motes

**Author notes:** Correspondence: Carlos Maluquer de Motes: Department of Microbial Sciences, University of Surrey, Guildford, GU2 7XH, United Kingdom,.

## Abstract

Cullin-RING E3 ubiquitin ligases (CRLs) have emerged as critical regulators of many cellular functions including innate immunity and inflammation. CRLs form multiprotein complexes in which specific adaptor proteins recruit the substrates to be ubiquitylated. Here, we systematically depleted all predicted SOCS-box proteins – the substrate adaptors for the CRL5 family - and assessed the impact on the activation of the NF-κB pathway. Depletion of SPSB1 resulted in a significant increase in NF-κB activation, indicating the importance of SPSB1 as an NF-κB negative regulator. In agreement, overexpression of SPSB1 suppressed NF-κB activity in a potent, dose-dependent manner in response to various agonists. Conversely, the activation of IRF-3, AP-1 and STATs was unaffected by SPSB1, showing its specificity for NF-κB. Mechanistically, SPSB1 suppressed NF-κB activation induced via multiple pathways including Toll-like receptors and RNA and DNA sensing adaptors, but was unable to prevent the phosphorylation and degradation of IκB nor the translocation of p65 into the nucleus. This indicated that SPSB1 exerts its inhibitory activity downstream, or at the level, of the NF-κB heterodimer and in agreement, SPSB1 was found to co-precipitate with p65. Additionally, A549 cells stably expressing SPSB1 presented lower cytokine levels including type I interferon in response to cytokine stimulation and virus infection. Taken together, our results reveal novel regulatory mechanisms in inflammation and innate immune signalling and identify the prominent role of SPSB1 in controlling NF-κB activation, thus providing new opportunities for the therapeutic targeting of NF-κB transcriptional activity.

## INTRODUCTION

Few transcription factors have such crucial roles in the induction of innate immune and inflammatory responses as the NF-κB family (1). NF-κB is central in the pathogenesis of multiple inflammatory disorders including those in the airway by inducing the production of pro-inflammatory cytokines such as interleukins (IL) and tumour necrosis factors (TNF). In addition, NF-κB contributes to the expression of type I interferon (IFN) in association with the IFN regulatory factors (IRF)-3/7, and the activator protein 1 (AP-1). Once secreted IFN triggers the production of hundreds of IFN-stimulated genes (ISG) via the Janus-associated kinase (JAK)-signal transducers and activators of transcription (STAT) signalling pathway, which confer an antiviral state to surrounding cells. NF-κB thus also impacts on the host antiviral response.

In the classical NF-κB pathway, the NF-κB p65 and p50 heterodimer is held inactive in the cytosol bound to the inhibitor of κB (IκB). Degradation of IκBα and subsequent release of NF-κB can be induced by multiple cytokine receptors and pattern-recognition receptors (PRRs) including Toll-like receptors (TLRs) that recognise viral and bacterial nucleic acids and lipids. Engagement of TNF-α with its receptor on the cell surface induces a signalling cascade that involves the TNFR-associated factor 2 (TRAF-2), whereas signalling downstream of TLRs and IL-1R employs TRAF-6. Activation of these signalling pathways induces the formation of polyubiquitin chains that act as a scaffold for the recruitment of the transforming growth factor (TGF)β-activated kinase 1 (TAK-1) complex via its TAK-1-binding proteins TAB2/3 and the IκB kinase (IKK) complex (2,3). Once both complexes are recruited, TAK-1 catalyses the phosphorylation and activation of the IKK catalytic subunits (IKK-α and IKK-β), which phosphorylate IκB (4). Phosphorylated IκBα is recognised by a Cullin-RING ubiquitin ligase (CRL) complex containing Cullin-1 and the F-box protein β-transducin repeat containing protein (β-TrCP), also known as FBXW11 (5,6), that mediates its ubiquitylation and proteasome-dependent degradation (7). The essential role of the β-TrCP-containing CRL1 complex in NF-κB activation is highlighted by the long list of viruses that antagonise its function including poxviruses (8,9), rotaviruses (10), and the human immunodeficiency virus (11).

Nuclear p65 associates with transcriptional activators such as p300/CBP and general transcription machinery and drives the expression of genes containing NF-κB responsive elements. A plethora of post-translational modifications (PTM) affecting p65 (i.e. phosphorylation, acetylation, methylation, ubiquitylation, etc.) modulate the potency of this response and its selectivity towards specific NF-κB-dependent genes (12). These modifications are critical in fine-tuning the transcriptional activity of NF-κB.

Ubiquitylation of a protein involves its covalent attachment of ubiquitin moieties that can subsequently be ubiquitylated to form ubiquitin chains (13). CRL are the largest family of ubiquitin ligases and are characterised by the presence of a Cullin (Cul) protein acting as a scaffold (14). There are 5 major Cul and hence CRL (CRL1-5) families, all of which share similar architecture but recruit and target different subset of substrates. CRL substrate recognition is directed by specific substrate receptor subunits. CRL5 complexes employ substrate receptor proteins containing the suppressor of cytokine signalling (SOCS) motif (15,16). The SOCS box in these adaptor proteins mediates interaction with the Elongin B/C proteins that associate with Cul-5, effectively forming a CRL5 complex also known as ECS (Elongin B/C-Cul5-SOCS-box protein).

Analogous to F-box proteins and CRL1 complexes, the SOCS-box domain appears in combination with other protein-protein interaction domains including Ankyrin, SPRY and Src homology domains (17,18). These domains are responsible for the recognition of substrates via unique signatures in the primary amino acid sequence or specific PTMs, but in many cases these remain to be elucidated. The most studied CRL5 complex is that formed by the so-called SOCS proteins (SOCS1-7), which target JAKs and act as potent inhibitors of JAK-STAT signalling (19). Here, we have performed an RNAi-based screen to assess the role of CRL5 adaptors in NF-κB signalling. Our work has identified several molecules positively and negatively regulating the pathway, in particular the SPRY and SOCS-box containing protein (SPSB)1. Depletion of SPSB1 resulted in enhanced NF-κB activation and cytokine expression, whereas its overexpression suppressed NF-κB responses triggered by cytokines as well as viruses. Our results indicate that SPSB1 associate with p65 but does not block its translocation, which suggests that it targets released p65. SPSB1 is known to target the inducible nitric oxide synthase (iNOS) (20,21) and has been linked with several pathways related to cancer, but no direct role for SPSB1 in controlling NF-κB activation has been reported. Our data therefore define another function for SPSB1 in innate immunity and inflammation and reveal novel regulatory mechanisms modulating NF-κB activation.

## RESULTS

### Identification of SPSB1 as a negative regulator of NF-κB signalling

In order to identify novel members of CRLs that act as regulators of the NF-κB pathway, we set up an assay to systematically deplete CRL genes in A549 cells stably expressing the firefly luciferase gene under the control of a synthetic NF-κB promoter (22). These cells were transfected with a library of small interfering (si)RNA commercially designed to target all predicted human SOCS-box containing proteins. Each gene was targeted by a pool of four individual siRNA duplexes. After 72 h the cells were treated with IL-1β and the luciferase activity was measured 6 h later. In a similar experiment, the viability of the cells transfected with the siRNA pools was measured. The screen was performed twice and included a non-targeting control (NTC) siRNA pool and a siRNA pool targeting β-TrCP, the CRL1 adaptor required for NF-κB activation. Each screen included triplicate replicates for each sample, and the data were normalised to the NTC (Fig. 1A). As expected knock-down of β-TrCP resulted in a decrease in NF-κB activation. Amongst the 38 siRNA pools tested only SPSB1 depletion consistently resulted in a significant increase (∼240%) in NF-κB activity without showing major changes in the viability of the cells (arbitrary cut-off of >75 %), suggesting that this protein has a prominent role in controlling NF-κB signalling.

**Figure 1.**
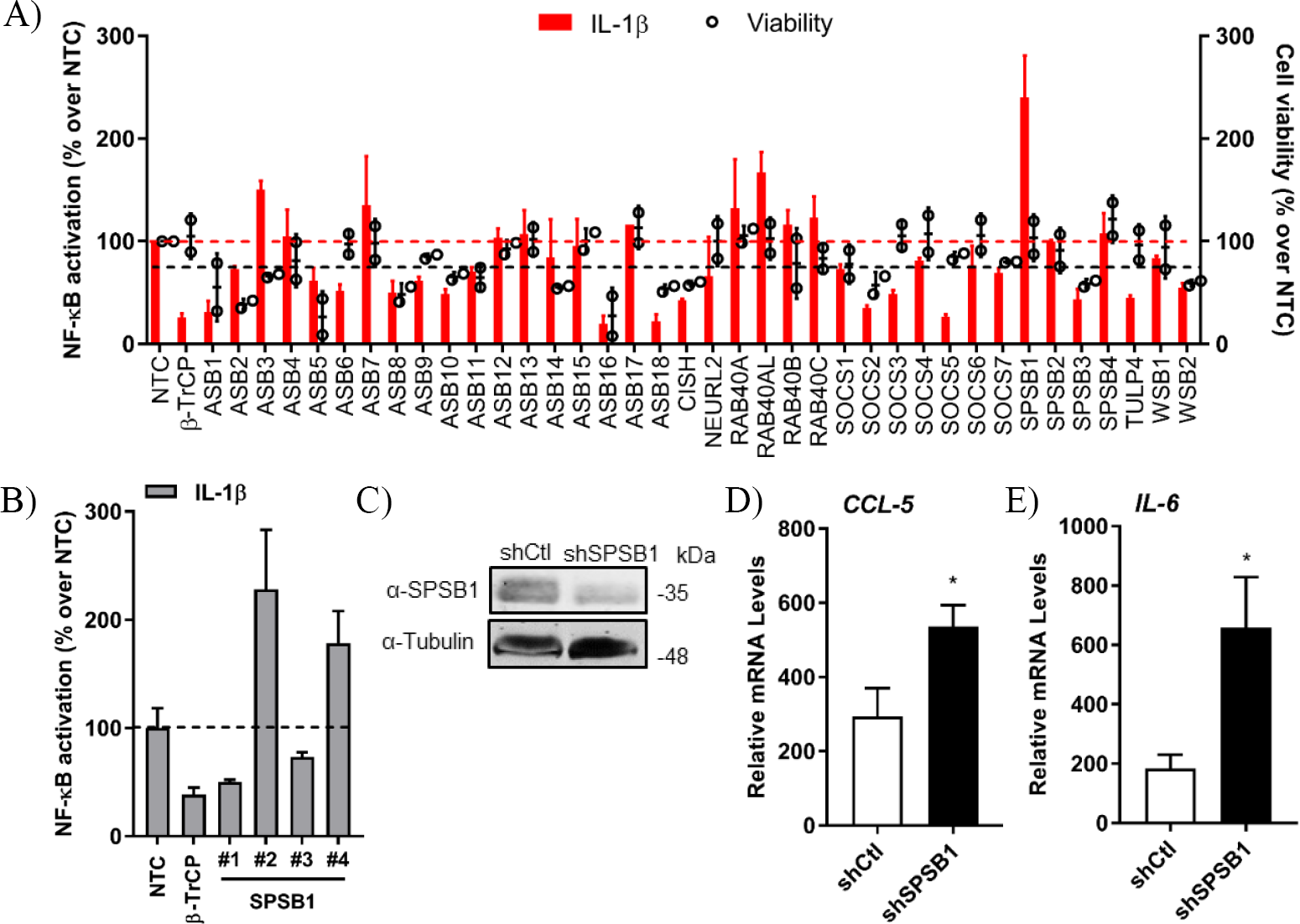
SPSB1 is a novel negative regulator of the NF-κB pathway. (A) IL-1β-mediated NF-κB activation (red bars, left axis) and cell viability (black circles, right axis) observed for A549-κB-Luc cells transfected with 30 nM siRNA targeting human CRL-5 adaptor proteins. Data were normalised to the non-stimulated condition and plotted as a % over NTC data. (B) The individual siRNA targeting SPSB1 from were tested individually and NF-κB activation was calculated as a % over NTC data. (C) A549 cells were transduced with a lentiviral construct containing shRNA against SPSB1 (shSPSB1) or NTC (shCtl). Cells were lysed in RIPA buffer and subjected to immunoblotting against endogenous SPSB1 and α-tubulin. (D-E) shSPSB1 and shCtl cells were treated with IL-1β (25 ng/ml) for 6 h, and the mRNA levels of (D) CCL-5 and (E) IL-6 were measured by qPCR. Means and standard deviations over the non-stimulated conditions are shown. Statistical significance was determined using an unpaired Student’s *t*-test (\**p* ≤ 0.05). In all panels, data are representative of at least 2 experiments performed independently and showing similar results.

To validate these initial data, we deconvolved the pool targeting SPSB1 and transfected the 4 different siRNA separately to test their effect on NF-kB activation under the same conditions used before, including NTC and β-TrCP siRNA controls. Two siRNA (#2 and #4) replicated the data observed for the pool (Fig. 1B) and this represented an H-score of 0.5, a value that supported the results from the first screen (23). We then performed stable depletion of SPSB1 via short hairpin (sh)RNA transduction. Depletion of SPSB1 in the shSPSB1 cells as compared to the NTC shCtl cells was confirmed by immunoblotting (Fig. 1C). These cell lines were then used to further confirm the impact of SPSB1 on NF-κB signalling. The cells were treated with IL-1β for 6 h and the mRNA levels of the cytokines *IL-6* and *CCL-5* were examined by quantitative PCR. Treatment with IL-1β resulted in 290-and 190-fold increase of *CCL-5* and of *IL-6* expression in the control A549 cell line, respectively. In the absence of SPSB1, this same treatment induced a significantly higher expression of both *CCL-5* and *IL-6* (540 and 660 fold, respectively) and this was statistically significant (Fig. 1D and 1E). Taken together, these data identified SPSB1 as a novel negative regulator of the NF-κB pathway, with its depletion resulting in higher expression of pro-inflammatory NF-κB-dependent genes.

### SPSB1 inhibits NF-κB activation, but not IRF-3, AP-1, or STAT activation

To study the function of SPSB1, its sequence was cloned into a mammalian expression vector containing 3 copies of the FLAG epitope at the N terminus. SPSB1 was then tested for its ability to inhibit NF-κB activation. HEK293T cells were transfected with a reporter expressing firefly luciferase under the control of the canonical NF-κB promoter, a control reporter expressing renilla luciferase, and either SPSB1 or the corresponding empty vector (EV). After 24 h, the NF-κB pathway was stimulated with IL-1β or TNF-α for a further 6 h. The ratio of firefly and renilla luciferase activities was calculated and plotted as a fold increase over the non-stimulated EV-transfected condition. The same cell lysates were also examined by immunoblotting to determine SPSB1 expression levels. Stimulation by IL-1β or TNF-α triggered >20-and >60-fold increase, respectively, in reporter activity in EV-transfected samples. Expression of SPSB1 reduced the activation induced by IL-1β (Fig. 2A) and TNF-α (Fig. 2B) in a dose-response and statistically significant manner.

**Figure 2.**
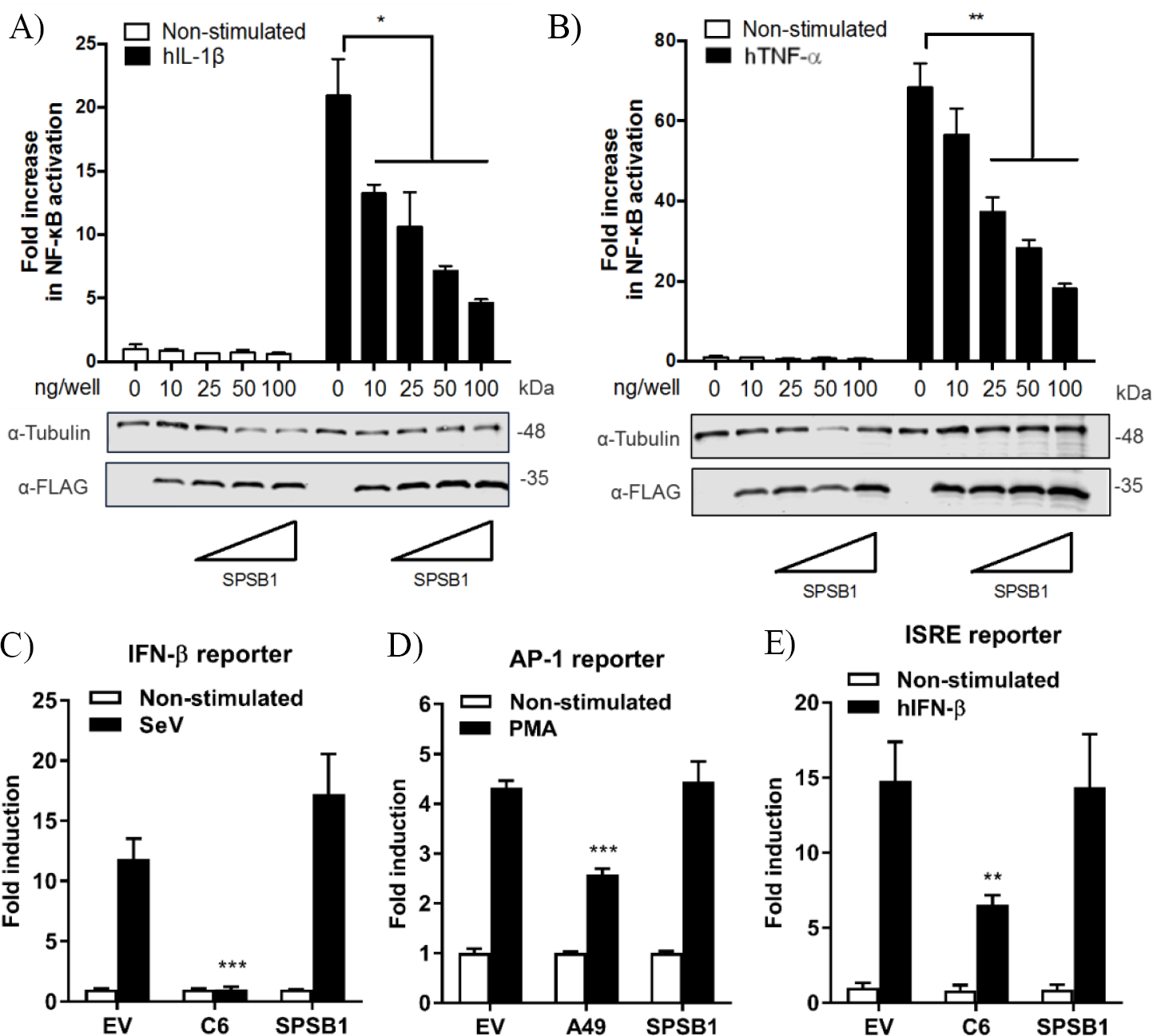
SPSB1 blocks NF-κB activation in response to IL-1β and TNF-α agonists, but it does not affect IRF-3, AP-1 or the JAK-STAT signalling pathway. (A-B) HEK293T cells were transfected with increasing doses of SPSB1-FLAG as indicated together with 70ng/well of pNF-κB-LUC and 10 ng/well of pTK-Renilla reporters. After 24 h the cells were stimulated with (A) 20 ng/mL of IL-1β or (B) 50 ng/mL of TNF-α for 6 h. (C-E) HEK293T cells were transfected with (C) 70 ng/mL of pIFNβ-LUC, (D) 200 ng/well of pAP1-LUC, or (D) 70 ng/well of pISRE-LUC as well as 10 ng/well pTK-Renilla reporters, together with 25 ng of SPSB1-FLAG, the indicated positive control plasmids, or the corresponding empty vector (EV). After 24 h the cells were stimulated with (C) SeV for a further 24 h, (D) 10 ng/ml of PMA for a further 24 h, or (E) 50ng/mL of IFN-β for a further 8 h. Data are representative of at least 3 independent experiments, each performed in triplicate. Means and standard deviations are shown and statistical significance was determined using an unpaired Student’s *t*-test (\**p* ≤ 0.05, \*\**p*≤ 0.01, \*\*\**p* ≤ 0.001).

To assess the specificity of SPSB1 in controlling innate immune responses, the activation of the IRF, mitogen-activated protein kinase (MAPK)/AP-1, and JAK-STAT signalling pathways was examined using reporter gene assays specific for each pathway. To determine the impact of SPSB1 on the IRF-3 signalling pathway, cells were transfected and subsequently infected with Sendai virus (SeV), a strong inducer of IRF-3 (24). The infection induced a 13-fold activation in the control cells. This was blocked by the vaccinia virus (VACV) protein C6, a known inhibitor of IRF-3 signalling (25), but remained unaffected by SPSB1 (Fig. 2C). Stimulation of the MAPK pathway was achieved by incubation with phorbol 12-myristate 13-acetate (PMA) for 24 h, which induced a 4-fold activation in the EV-transfected cells. This response was downregulated by the VACV protein A49 (26), but not by SPSB1 (Fig. 2D). Finally, to address whether SPSB1 was able to impact signalling triggered by IFN via the JAK/STAT pathway, cells were transfected with a reporter expressing luciferase under the control of the IFN stimulating response element (ISRE) and stimulated with IFN-β. This treatment resulted in a 15-fold induction in both EV-as well as SPSB1-transfected cells, indicating that SPSB1 did not affect JAK/STAT signalling (Fig. 2E). The above demonstrates that SPSB1 is not a general transcriptional modulator and specifically regulates NF-κB responses.

### SPSB1 inhibits NF-κB activation downstream of multiple effectors

We then aimed to gain further insights into SPSB1 regulation of NF-κB signalling using 2 different approaches. First, we performed luminescence-based mammalian interactome mapping (LUMIER) assays (27–29) to examine possible interactions between SPSB1 and a number of molecules operating at multiple levels downstream of the IL-1R signalling cascade (Fig. 3A). SPSB1 was initially found to self-associate (Fig. S1). This property was used to ensure that the fusion of SPSB1 with renilla luciferase (Rluc) did not affect its expression and ability to interact with FLAG-SPSB1 (Fig. S2). FLAG-SPSB1 was then co-transfected with Rluc fusions for either TRAF-6, TAK-1, TAB-2, TAB-3, IKK-α, IKK-β, IKK-γ and β-TrCP as well as SPSB1 or an Rluc only construct. Rluc activity was measured before and after immunoprecipitation of FLAG-SPSB1 and RLuc ratios were calculated. Using this assay none of the tested NF-κB components interacted with SPSB1 (Fig. S1C). The second approach relied on reporter assays in which NF-κB was triggered by over-expression of signalling molecules. SPSB1 was able to suppress NF-κB activation deriving from the adaptors TRAF-6 (Fig. 3B) and TRAF-2 (Fig. 3C); the RNA sensor RIG-I (Fig. 3D), which activates NF-kB at the level of TRAF-6; the kinase IKK-β (Fig. 3E); and the DNA sensors cGAS and STING (Fig. 3F), which converge on the NF-κB pathway at the level of the IKK complex. Taken together, these data indicated that SPSB1 acted downstream of these molecules. In agreement with this observation SPSB1 inhibited NF-κB activation triggered by p65 over-expression (Fig. 3G).

**Figure 3.**
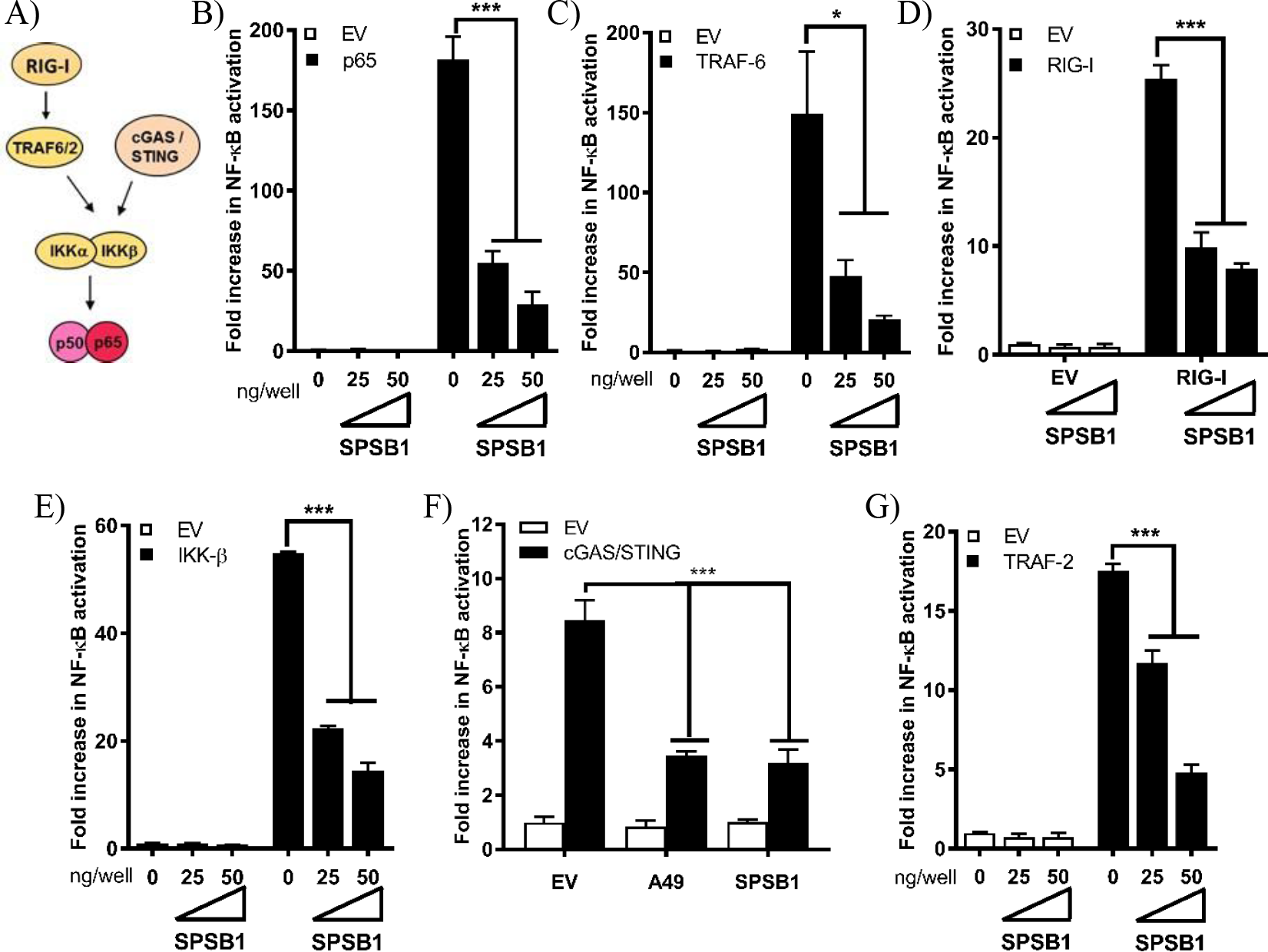
SPSB1 blocks NF-κB signalling downstream, or at the level, of the p50/p65 complex. (A) Schematic representation of the NF-κB key molecules and the cross-link with the cGAS/STING pathway. The arrows indicate the sequence of activation. (B-G) HEK293T cells were transfected with 70 ng/well of pNF-κB-LUC and 10 ng/well of pTK-Renilla, and two doses of SPSB1-FLAG (25 and 50 ng) or an empty vector (EV). The cells were stimulated by co-transfection of (B) 5 ng/well of TRAF-6, (C) 10 ng/well of TRAF-2, (D) 5 ng/well of RIG-I-CARD, (E) 50 ng/well of IKK-β, (F) 20 ng/well of cGAS and 20 ng/well of STING, or (G) 2 ng/well of p65. VACV protein A49 was used as a control. Data are representative of at least 3 independent experiments, each performed in triplicate. Means and standard deviations are shown and statistical significance was determined using an unpaired Student’s *t*-test (\**p* ≤ 0.05, \*\*\**p* ≤ 0.001).

### SPSB1 does not interfere with IκBα phosphorylation or degradation

If SPSB1 blocked p65-induced NF-κB activation, IκBα should be phosphorylated and degraded in the presence of SPSB1. To verify this, we first generated A549 stable cell lines expressing SPSB1 or GFP as a control protein using lentiviruses. Immunoblotting against FLAG revealed the successful transduction and expression of SPSB1 (Fig. 4A). To validate that SPSB1 expression was sufficient and functional in these cells, the expression of *CCL-5* (Fig. 4B), *ICAM-1* (Fig. 4C) and *iNOS* (Fig. 4D), all of which contain NF-κB sites in their promoters, was assessed by quantitative PCR in response to IL-1β stimulation. The expression of all these genes was enhanced upon IL-1β treatment, but to a lower extent in cells expressing SPSB1.

**Figure 4.**
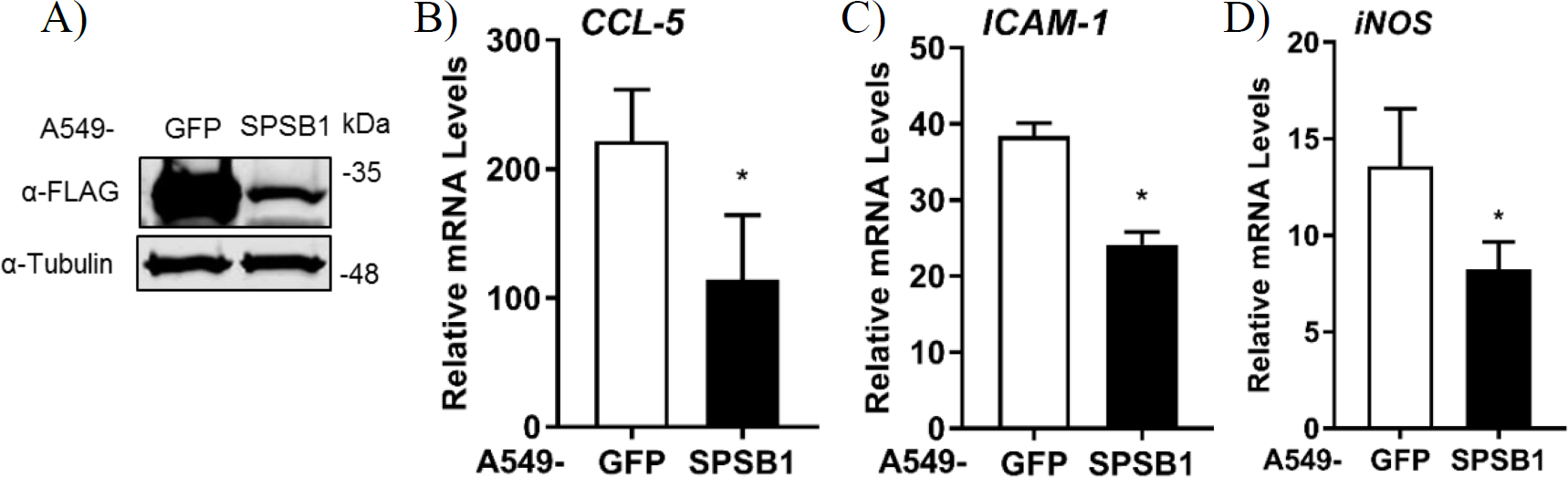
Expression levels of NF-κB-dependent genes in A549 cells stably expressing SPSB1 upon induction with IL-1β. (A) A549 cells were transduced with lentiviral constructs expressing FLAG-GFP or SPSB1. Cells were harvested in RIPA buffer and subjected to immunoblotting against FLAG and α-tubulin. (B-D) A549 cells stably expressing GFP or SPSB1 were treated with 25 ng/ml of IL-1β for 6 h. The mRNA levels of (B) CCL-5, (C) ICAM-1, and (D) iNOS were measured by qPCR. Data are presented as fold increase over the non-stimulated condition and representative of at least 2 independent experiments, each performed in triplicate. Means and standard deviations are shown and statistical significance was determined using an unpaired Student’s *t*-test (\**p* ≤ 0.05).

We then assessed the kinetics of phosphorylation and degradation of IκBα in these cells. Exposure to IL-1β induced significant p-IκBα levels in as little as 5 mins and this was concomitant with subsequent degradation of IκBα (Fig. 5). The presence of SPSB1 had no effect on either the intensity or the kinetics of phosphorylation of IκBα, nor its subsequent degradation. We also assessed the phosphorylation of Ser536 in p65, a cytoplasmic event that relates to p65 activation (30). No differences in p-p65 levels were observed between SPSB1 and its control cell line (Fig. 5). In addition, the total levels of p65 remained similar in the presence of SPSB1, suggesting that this protein does not affect p65 turnover.

**Figure 5.**
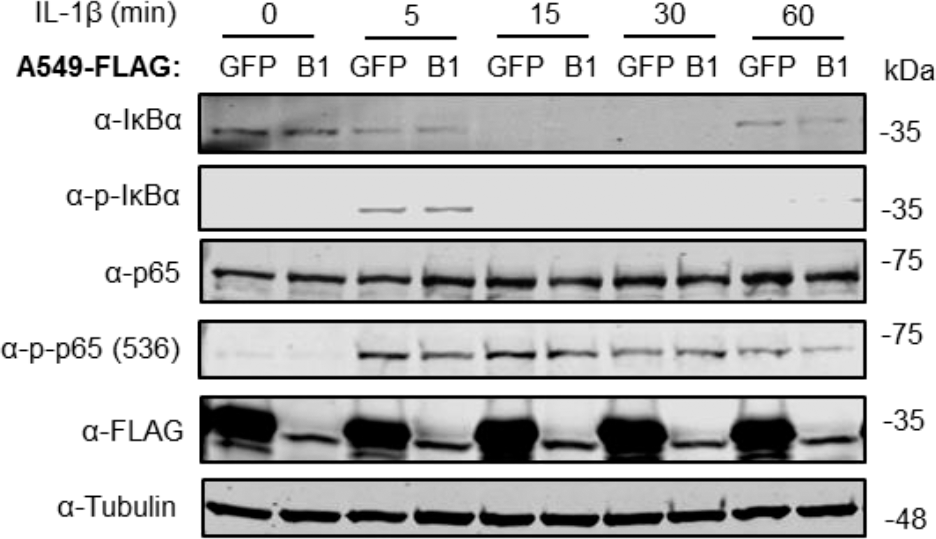
IκBα degradation in the presence of SPSB1. A549 cells stably expressing FLAG-GFP or FLAG-SPSB1 were treated with 25 ng/ml of IL-1β for the indicated length of time. Cells were harvested in RIPA buffer supplemented with protease and phosphatase inhibitors and lysates were subjected to immunoblotting against the indicated proteins. Data are representative of 3 experiments performed independently.

### SPSB1 associates with p65 but does not interfere with its translocation

When IκBα is phosphorylated and degraded, the NF-κB heterodimer is free to translocate into the nucleus and induce the expression of NF-κB-dependent genes. We therefore assessed whether p65 would translocate in the presence of SPSB1. Cells were challenged with IL-1β and after 30 mins stained for p65 and SPSB1 (FLAG). In control unstimulated cells p65 located in the cytosol and moved to the nucleus upon stimulation (Fig. 6A). In SPSB1-expressing cells, p65 translocated to the nucleus to the same extent upon IL-1β exposure and no differences were observed. This indicated that SPSB1 was not able to restrict p65 translocation. Interestingly, SPSB1 showed both nuclear and cytosolic distribution in unstimulated cells, but a notable nuclear localisation after stimulation, indicating that either SPSB1, or its target, alters its cellular distribution in response to NF-κB signalling. The fact that SPSB1 shadows p65 nuclear translocation and inhibits p65-induced NF-κB activation (Fig. 3G) suggested an association between SPSB1 and p65. We thus immunoprecipitated HA-tagged p65 in the presence of SPSB1 or GFP as a control and observed a specific co-precipitation of SPSB1 with p65 (Fig. 6B). Collectively, these data revealed that SPSB1 is a potent suppressor of NF-κB responses that interacts with p65 in a manner that does not affect translocation.

**Figure 6.**
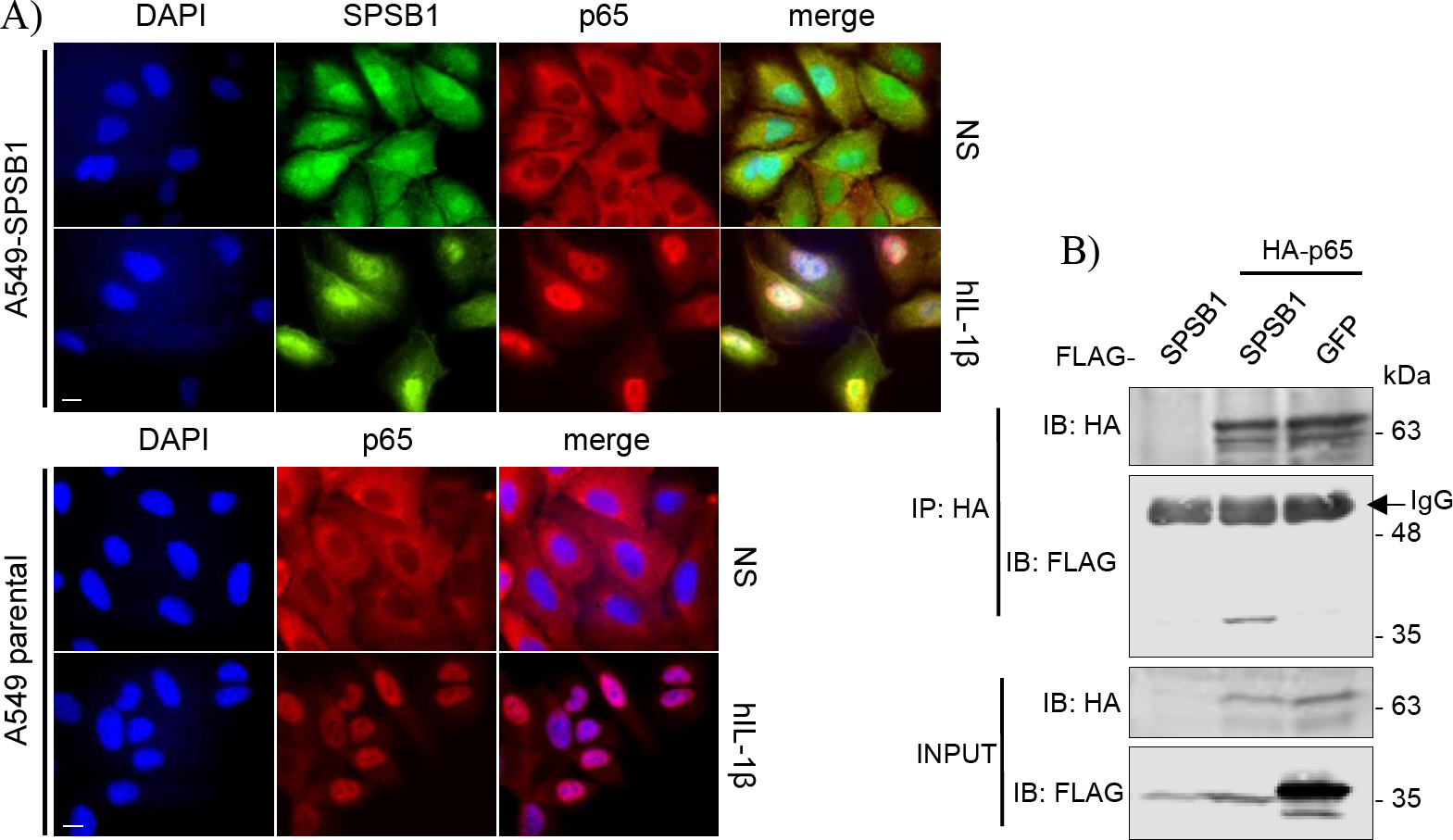
SPSB1 associates with p65, but does not interfere with its translocation. (A) A549 cells stably expressing FLAG-SPSB1 or their parental control were treated with 25 ng/mL of IL-1β for 30 mins or left unstimulated (NS). The cells were then fixed and stained for FLAG (green) and p65 (red) together with DAPI (blue) to visualise cell nuclei. Merged images are also shown. Images are representative of multiple visual fields showing the same results. Bar, 10 μm. (B) HEK293T cells transfected with HA-p65 and FLAG-SPSB1 or GFP were lysed and subjected to HA immunoprecipitation (IP). Lysates and final IP eluates were immunoblotted (IB) against the indicated tags. IgG heavy chain is indicated with an arrow.

### SPSB1 inhibits NF-κB activation induced by RSV infection

Viruses are common inducers of NF-κB signalling. Respiratory syncytial virus (RSV) is a common respiratory pathogen and the main cause of airway inflammation in infants, and it is known to trigger NF-κB and type I IFN responses in the airway (31). To address the role of SPSB1 in controlling virus-induced responses, we infected our A549 cell lines with 2 PFU/cell of RSV and performed qPCR analysis on a number of cytokines. RSV infection induced a high expression of *CCL-5*, higher than that observed upon IL-1β treatment, and this was reduced in the presence of SPSB1 in a statistically-significant manner (Fig. 7A). Similarly, RSV triggered measurable levels of *IFN-β* in these cells and this was reduced SPSB1 (Fig. 7B). Interestingly, we also observed a significant reduction in the levels of IFN-dependent genes such as *ISG54* and *OAS1* (Fig. 7C and 7D). Given the inability of SPSB1 to directly downregulate the JAK-STAT signalling pathway, these results indicate that SPSB1 effective suppression of IFN-β production impacted on the expression of antiviral genes.

**Figure 7.**
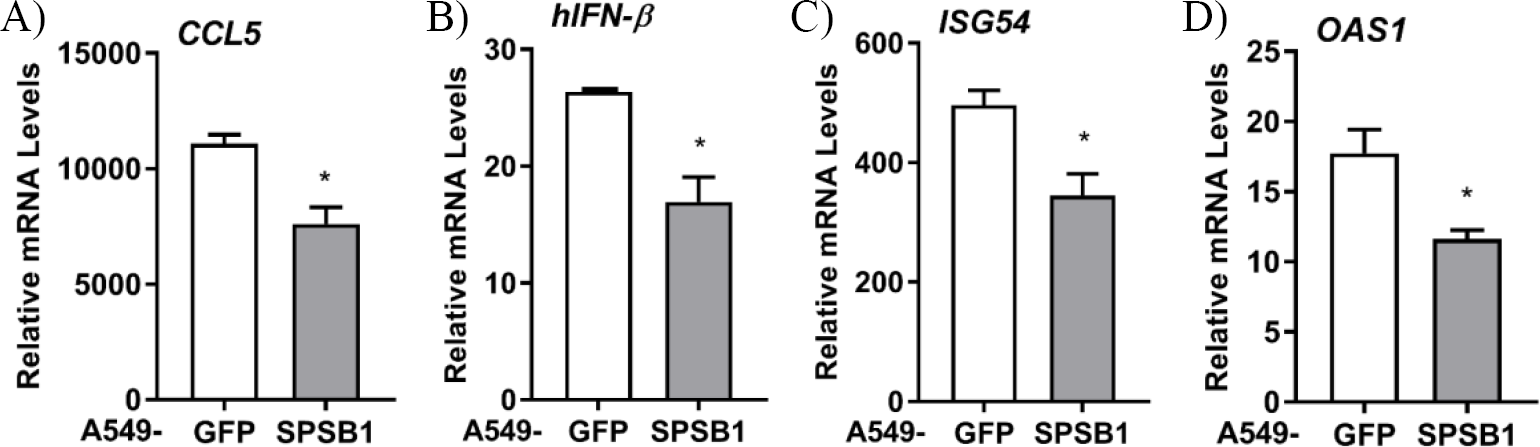
SPSB1 inhibits the innate immune response to RSV infection. A549 cells stably expressing GFP or SPSB1 were infected with 2 PFU/cell of RSV for 24 h. The mRNA levels of (A) CCL5, (B) IFN-β, (C) ISG54 and (D) OAS1 were measured by qPCR. Data are representative of at least 3 independent experiments, each performed in triplicate. Means and standard deviations are shown and statistical significance was determined using an unpaired Student’s *t*-test (\**p* ≤ 0.05).

## DISCUSSION

Here we have explored the role of CRL5 complexes in NF-κB activation in airway epithelial cells using an unbiased screen for SOCS-box proteins. This screen has identified SPSB1 as a novel regulator of the pathway: SPSB1 depletion resulted in enhanced NF-κB-dependent cytokine production (Fig. 1) and this effect was reversed by its overexpression (Fig. 2, 4 and 7). In addition, our results indicate that SPSB1 controls NF-κB activation when cells were exposed to inflammatory cytokines (Fig. 4) as well as viruses (Fig. 7). Therefore, our work highlights SPSB1 as a novel and important participant in the signalling network that governs the production of NF-κB-dependent cytokines.

SPSB1 is the first member of the SPSB family, a group of 4 proteins (SPSB1-4) characterised by the presence of a SPRY domain and a C-terminal SOCS box domain that engages with the CRL E3 ubiquitin ligase complex (32). SPSB1, SPSB2 and SPSB4 are known to target the inducible nitric oxide synthase (iNOS) via the SPRY domain and trigger its proteasomal degradation (20,21,33). In addition, SPSB1 regulates multiple cancer-associated pathways via interactions with c-met (34,35), the apoptosis-related protein Par-4 (36,37) and the TGF-β receptor (38,39). The specific motif that SPSB1 recognises on its targets was suggested to be (D/E)-(I/L)-N-N-N. However, the degron recognised by SPSB1 in the TGF-β receptor has been mapped to N-I-N-H-N-T (39). The difference in sequence of the proposed motifs suggests the existence of more SPSB1 degrons than previously inventoried. Interestingly, SPSB1 has recently been shown to direct non-degradative ubiquitylation in the nucleus to regulate alternative splicing (40). Our results reveal that SPSB1 restricts the extent of NF-κB activation induced by cytokines and viruses downstream of IκBα degradation and p65 translocation and associates with p65. This indicates that SPSB1 targets p65 in the nucleus or in the cytosol in a manner that does not affect its ability to translocate. SPSB1 targeting affects the transactivation potential of p65, but not its stability. An interesting possibility is that SPSB1 mediates non-degradative ubiquitylation of p65 itself and affects its transcriptional activity perhaps by competing with other PTM known to activate p65 such as acetylation (41). Thus, SPSB1 would limit signal-induced p65 activation without triggering ubiquitin-dependent proteolysis. The ability of SPSB1 to catalyse non-degradative K29 ubiquitin chains has already been described (40).

An SPSB1 substrate that is of particular importance in innate immunity and inflammation is iNOS (also known as NOS2) and its catalytic product NO. iNOS is an inducible gene that is expressed at low levels in human respiratory epithelia and is upregulated in disease. Activation of NF-κB and STAT1 in response to TLR agonists and cytokines is largely responsible for the transcriptional induction of iNOS expression (42). Interestingly, SPSB1 expression is also enhanced by NF-κB and type I IFN (20). This indicates that SPSB1 expression is tightly regulated and that, according to the data presented here, represents a negative feed-back loop on NF-κB. This also implies that SPSB1 has a dual role controlling iNOS: i) it limits iNOS expression by downregulating NF-κB activation, and ii) it drives iNOS ubiquitylation and its proteasome-dependent destruction. This indicates that SPSB1 is a unique molecule controlling inflammatory responses. We did not observe a role for other iNOS-modulating CLR5 complexes such as SPSB2 or SPSB4 in inhibiting NF-kB activation, although we cannot rule out that the RNAi depletion for SPSB2 and SPSB4 in our screen was not sufficiently efficient. It would therefore be interesting to assess the role of these molecules as well as its paralogue FBXO45 in the regulation of NF-κB signalling.

Our screen has also highlighted other molecules that might regulate NF-κB signalling. For instance, depletion of SOCS3 or SOCS5 led to a substantial reduction in NF-κB activation. SOCS3 was originally identified as a negative regulator of the JAK/STAT pathway (43) by binding to specific JAK complexes and preventing downstream signal transduction (44). Additionally, it was shown to inhibit NF-κB activity (45). Here, RNAi against SOCS3 resulted in low NF-κB activation (48 % upon stimulation), suggesting that SOCS3 is a positive regulator of this pathway. This apparent contradiction may point out to cell-specific role for SOCS3. SOCS5 has also been identified as a positive regulator of NF-κB signalling in our screen. SOCS5 has been shown to regulate IL-4 (46) and epidermal growth factor receptor EGFR) signalling (47,48). In addition, inhibition of EGFR/PI3K signalling by SOCS5 conferred protection against influenza infection (49). Our data suggest that SOCS5 is a critical factor required for NF-κB activation since its depletion severely impaired NF-κB reporter activation (26 % upon stimulation). This may indicate that SOCS5 is necessary to activate NF-κB upon influenza virus infection and mount a protective response. Finally, we also noticed that depletion of SOCS1, a known inhibitor of NF-κB and JAK/STAT signalling (50,51), did not result in significant changes in NF-κB activation in our screen. Further analysis of gene expression data revealed that SOCS1 is not expressed in A549 cells (52), which accounts for these results.

Excessive inflammation is central to a large number of pathologies. For instance, in the respiratory tract obstructive lung diseases such as asthma or chronic obstructive pulmonary disease (COPD) are characterised by inflammatory gene expression and the production of inflammatory mediators that enhance the recruitment of inflammatory cells (53). NF-κB is an important player in this multifactorial diseases as evidenced by the fact that the therapeutic efficacy of the main treatment for asthma - glucocorticoids – is thought to be largely caused by their ability to suppress NF-κB and AP-1 responses (54). In these diseases, NF-κB occurs largely in response to cytokines such as IL-1β and TNF-α or by infection with viruses during exacerbations (55). We present here the novel finding that SPSB1, a member of the CRL5 family, downregulates the expression of inflammatory cytokines and other NF-κB-dependent genes in airway epithelial cells exposed to cytokines and viral infection. SPSB1 may thus have regulatory functions in chronic inflammatory disorders of the respiratory tract as well as acute virus-induced exacerbations. Our work reveals a new connection between CRL and innate immunity and may offer alternative strategies for the manipulation of NF-κB transcriptional activity in inflammatory pathologies.

## EXPERIMENTAL PROCEDURES

### Cells, viruses and agonists

A549 and HEK293T cells were grown in Dulbecco Modified Eagle medium (DMEM, Life Technologies) supplemented with 10 % heat-inactivated fetal calf serum (FCS, Biowest), 100 U/mL penicillin and 100 μg/mL Streptomycin (Pen/Strep, Invitrogen). The A549-κB-LUC cells were previously described (22). A549-SPSB1 and A549-GFP stable cell lines were grown as above with the addition of 2 μg/mL puromycin (Invitrogen). RSV strain A2 was from Gill Elliott (University of Surrey, United Kingdom). IL-1β, TNF-α and IFN-β were from Peprotech. PMA was from Santa Cruz.

### RNAi depletion screens

siRNA sequences targeting CRL5 adaptors were purchased from Horizon Discovery and resuspended in nuclease-free water to 1 μM final concentration. A549-κB-LUC cells were reverse-transfected in triplicate replicates with 30 nM siRNA using Interferin-HTS (Polyplus) and incubated for 72 h. The cells were then stimulated with 1 ng/mL of IL-1β for 6 h and subsequently washed with ice-cold PBS and lysed in Passive Lysis Buffer (PLB; Promega). Luciferase activity was measured in a Clariostar plate reader (BMG Biotech) and data for each sample were normalised to its non-stimulated condition and plotted as mean % ± SD over the NTC-transfected control. Data shown are representative of 2 independent screens showing similar results. Cells were also reverse-transfected in identical manner to determine cell viability 72 h post-transfection using CellTiter-Glo (Promega) following manufacturer’s recommendations.

### shRNA depletion and overexpression of SPSB1 in A549 cells

A549 cells were depleted for SPSB1 using specific short hairpin sequences expressed from the HIV-1-based shRNA expression vector HIVSiren (56). These shRNA sequences were generated using an open-access algorithm (RNAi designer, ThermoFisher) and were as follows: Top shRNA strand: 5’-GATCCACACAACCCTCGTGGGGAACGAATTCCCCACGAGGGTTGTG-3’; bottom SPSB1 strand: 5’-AATTCACAACCCTCGTGGGGAATTCGTTCCCCACGAGGGTTGTGTG-3’. Lentiviral particles were produced by transfection of HEK293T cells (seeded in 100 mm dishes) with 9 μg of pHIVSIREN shRNA, 5 μg of p8.91 packaging plasmid (57), and 3 μg of vesicular stomatitis virus-G glycoprotein expressing plasmid pMDG (Genscript) using polyethylenimine (PEI; Polysciences). Virus supernatants were harvested at 48 and 72 h post-transfection, pooled, and used to transduce A549 cells, which were subsequently selected for puromycin resistance (Invivogen, 2 μg/ml).

To generate A549 cells overexpressing SPSB1, SPSB1 was PCR amplified using primers 5’-GAA GCG GCC GCG GGT CAG AAG GTC ACT GAG-3’ (fwd) and 5’-GAC TCT AGA TCA CTG GTA GAG GAG GTA GG-3’ (rev). The PCR product was subsequently ligated into a pcDNA4/TO expression vector (Invitrogen) previously modified to express genes in frame with 3 N-terminal copies of the FLAG epitope, an N-terminal copy of the V5 epitope, or an N-terminal tandem affinity purification tag containing 2 copies of the strep tag and 1 copy of the FLAG tag as previously described (29,58). FLAG-SPSB1 was then subcloned into a lentivirus vector carrying puromycin resistance (gift from Greg Towers, University College London). Lentiviral particles were produced in HEK293T as above and virus supernatants were harvested at 48 and 72 h post-transfection.

### Quantitative PCR

RNA from confluent 6-well plates of A549 cells was purified using the Total RNA Purification Kit (Norgen Biotech). One μg of RNA was transformed into cDNA using Superscript III reverse transcriptase (Invitrogen). cDNA was diluted 1:5 in water and used as a template for real-time PCR using SYBR® Green PCR master mix (Applied Biosystems) in a LightCycler® 96 (Roche). Expression of each gene was normalized to an internal control (18S) and these values were then normalized to the shCtl or GFP control cells to yield a fold induction. Primers used for the detection of CXCL10 (59), CCL5 (8), IFNβ and 18S (60) have been described. Data shown are representative of at least 3 independent experiments showing similar results, each performed in triplicate and plotted as mean ± SD.

### Reporter gene assays

HEK293T cells were seeded in 96-well plates and transfected with the indicated reporters and expression vectors using PEI as described in the figure legends. The reporter plasmids have been described previously (8). After 24 h the cells were stimulated by exposure to different agonists or by co-transfection with activating plasmids as indicated in the figure legends. Plasmids for signalling molecules have been described (58) with the exception of untagged and HA-tagged p65 that were from Geoffrey Smith (University of Cambridge, United Kingdom). TAP-tagged VACV C6 (25) and HA-tagged VACV A49 (8) have been described. After stimulation cells were washed with ice-cold PBS and lysed with PLB. Luciferase activity was measured in a Clariostar plate reader and firefly and renilla ratios were calculated for each condition. Data were normalised to mock-infected samples or samples transfected with an empty vector and presented as a fold increase. In all cases data shown are representative of at least 3 independent experiments showing similar results, each performed in triplicate and plotted as mean ± SD.

### LUMIER assays

HEK293T cells were co-transfected with FLAG-SPSB1 and Rluc fusions for NF-κB signalling components for 24 h. Rluc fusions were from Felix Randow and/or have been previously described (28,61–63). Cells were lysed in IP buffer (20mM Tris-HCl pH7.4, 150mM NaCl, 10mM CaCl2, 0.1 % Triton-X and 10 % glycerol) supplemented with protease inhibitors (Roche) and cleared lysates were subjected to affinity purification (AP) with streptavidin beads for 6 h at 4 °C. The beads were then washed 3 times with lysis buffer prior to elution with biotin (10 mg/ml) diluted in PLB. Luciferase activity was measured and data were plotted as a binding fold over Rluc-only control.

### Immunoprecipitation

HEK293T cells were seeded in 10-cm dishes and transfected with 5 μg of the indicated plasmids using PEI. After 24 h cells were lysed with IP buffer as above. Cleared lysates were incubated with HA antibody (Sigma) for 16 h at 4 °C and subsequently Protein G beads (Santa Cruz) were added for a further 2 h. The beads were then washed 3 times with IP buffer prior to incubation at 95 °C for 5 min in Laemmli loading buffer to elute bound proteins. Cleared lysates and FLAG eluates were analysed by SDS-PAGE and immunoblotting. Data shown are representative of at least 3 independent experiments showing similar results.

### SDS-PAGE and immunoblotting

Cells were lysed in IP or RIPA buffer (50mM Tris-HCl pH8, 150mM NaCl, 1 % NP-40, 0.5 % sodium deoxycholate and 0.1 % SDS) and resolved by SDS-PAGE and transferred to nitrocellulose membranes (GE Healthcare) using a Trans-Blot® semi-dry transfer unit (Bio-Rad). Membranes were blocked in 0.1 % Tween PBS supplemented with 5% skimmed milk (Sigma) and subjected to immunoblotting with the following primary antibodies at the indicated dilutions: SPSB1 (Abcam, 1:1,000); FLAG (Sigma, 1:1,000); IκBα (Cell Signalling, 1:1,000); p-IκBα (S32/36) (5A5) (Cell Signalling, 1:1,000); p65 (Santa Cruz, 1:500); p-p65 (Ser536) (Santa Cruz, 1:500); V5 (BioRad, 1:5,000); α-Tubulin (Upstate Biotech, 1:10,000); HA (Sigma, 1:2,000). Primary antibodies were detected using IRDye-conjugated secondary antibodies in an Odyssey Infrared Imager (LI-COR Biosciences).

### Immunofluorescence

Cells were seeded into 6-well plates containing sterile glass coverslips. Following stimulation with IL-1β (25 ng/ml) for 30 min, the cells were washed twice with ice-cold PBS and fixed in 4 % (w/v) paraformaldehyde. The cells were then quenched in 150 mM ammonium chloride, permeabilized in 0.1 % (v/v) Triton X-100 in PBS, and blocked for 30 min in 5 % (v/v) FBS in PBS. The cells were stained with rabbit anti-FLAG (Sigma, 1:300) and mouse anti-p65 antibody (Santa Cruz, 1:50) for 1 h, followed by incubation with goat anti-rabbit IgG Alexa Fluor 488 and goat anti-mouse IgG Alexa Fluor 568 secondary antibodies (Invitrogen). Coverslips were mounted in Mowiol 4-88 (Calbiochem) containing DAPI (4′,6-diamidino-2-phenylindole). Images were taken on a LSM 510 META confocal laser scanning microscope (Zeiss) using the LSM image browser software (Zeiss).

### Statistical analysis

Statistical significance was determined using an unpaired Student’s *t*-test with Welch’s correction where appropriate.

## ACKNOWLEDGEMENTS

The authors would like to thank GJ Towers, G Elliott, F Randow and GL Smith for providing reagents. This work was supported by an Asthma UK Innovation Grant (AUKIG2016353) to CMdM. IG was supported by a University of Surrey School of Biosciences and Medicine studentship.

---

**Figure S1.**
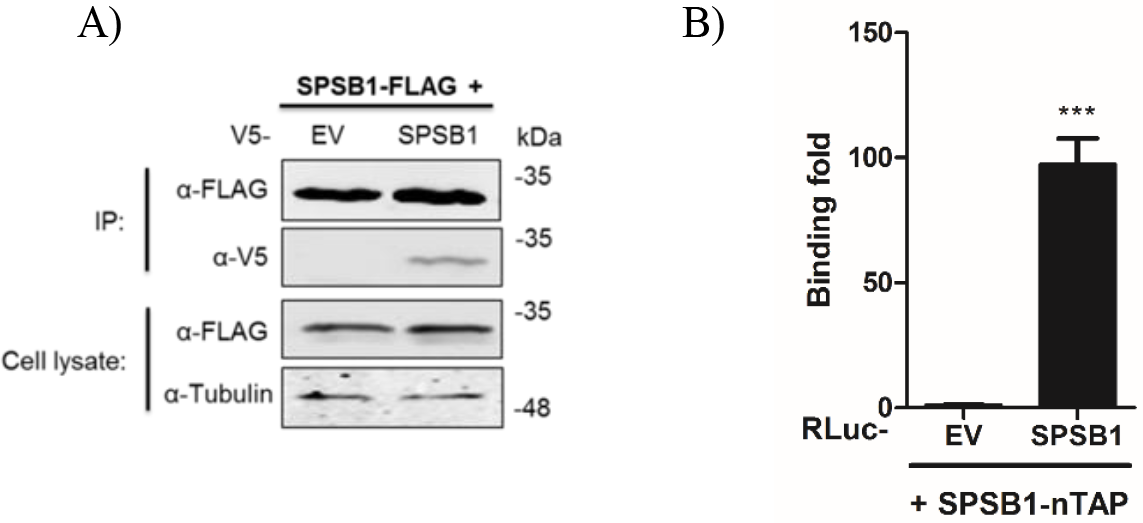
SPSB1 self-associates. (A) HEK293T cells were co-transfected with V5-tagged SPSB1, or an empty vector (EV) and FLAG-tagged SPSB1. The cells were lysed in IP buffer and subjected to FLAG IP. The IP and cell lysates were loaded onto an SDS-PAGE gel and immunoblotted against V5, FLAG and α-tubulin. (B) HEK293T cells were co-transfected in triplicates with RLuc-tagged SPSB1, or RLuc-only (EV) as a control and SPSB1-nTAP. Cells were lysed in IP buffer supplemented with protease inhibitors and lysates subjected to AP using streptavidin beads. The beads were eluted with biotin and the luciferase activity was measured both in lysates and eluates. The binding fold was calculated as an eluate/lysate ratio and this was normalised to the EV control. The experiments have been repeated three times independently. Means and standard deviations are shown and statistical significance was determined by using an unpaired Student’s *t*-test (\*\*\**p* ≤ 0.001).

**Figure S2.**
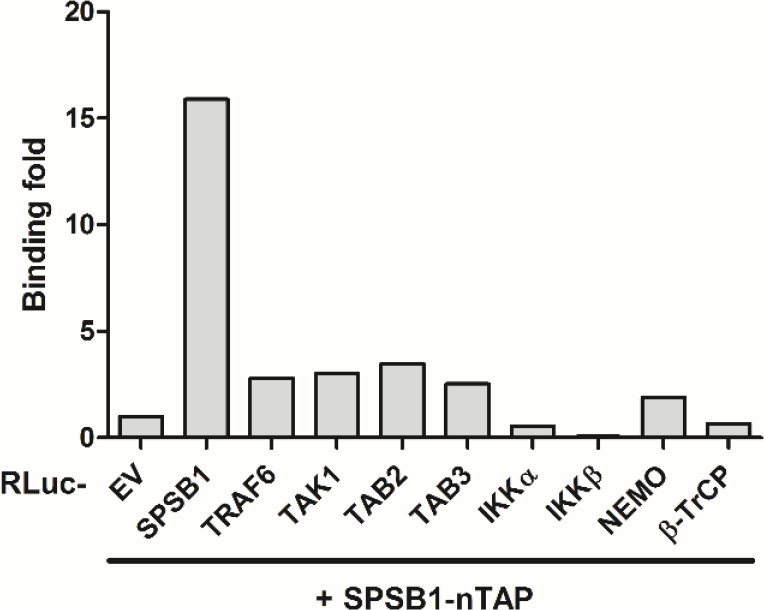
SPSB1 does not associate with any of the NF-κB components tested in LUMIER screens. HEK293T cells were co-transfected with SPSB1-nTAP and different components of the NF-κB pathway fused to RLuc, or an Rluc-only (EV) control. SPSB1 self-association was used a positive control. Cells were treated as in S1 and the binding fold was calculated as an eluate/lysate ratio and this was normalised to the EV control. Data are representative of 2 experiments showing similar results.

## References

1. Ghosh, S., and Hayden, M. S. (2012) Celebrating 25 years of NF-kappaB research. Immunol Rev 246, 5–13

2. Chen, Z. J. (2012) Ubiquitination in signaling to and activation of IKK. Immunol Rev 246, 95–106

3. Clark, K., Nanda, S., and Cohen, P. (2013) Molecular control of the NEMO family of ubiquitin-binding proteins. Nat Rev Mol Cell Biol 14, 673–685

4. Traenckner, E. B., Pahl, H. L., Henkel, T., Schmidt, K. N., Wilk, S., and Baeuerle, P. A. (1995) Phosphorylation of human I kappa B-alpha on serines 32 and 36 controls I kappa B-alpha proteolysis and NF-kappa B activation in response to diverse stimuli. EMBO Journal 14, 2876–2883

5. Cenciarelli, C., Chiaur, D. S., Guardavaccaro, D., Parks, W., Vidal, M., and Pagano, M. (1999) Identification of a family of human F-box proteins. Curr Biol 9, 1177–1179

6. Winston, J. T., Koepp, D. M., Zhu, C., Elledge, S. J., and Harper, J. W. (1999) A family of mammalian F-box proteins. Curr Biol 9, 1180–1182

7. Yaron, A., Hatzubai, A., Davis, M., Lavon, I., Amit, S., Manning, A. M., Andersen, J. S., Mann, M., Mercurio, F., and Ben-Neriah, Y. (1998) Identification of the receptor component of the IkappaBalpha-ubiquitin ligase. Nature 396, 590–594

8. Mansur, D. S., Maluquer de Motes, C., Unterholzner, L., Sumner, R. P., Ferguson, B. J., Ren, H., Strnadova, P., Bowie, A. G., and Smith, G. L. (2013) Poxvirus targeting of E3 ligase beta-TrCP by molecular mimicry: a mechanism to inhibit NF-kappaB activation and promote immune evasion and virulence. PLoS Pathog 9, e1003183

9. Neidel, S., Maluquer de Motes, C., Mansur, D. S., Strnadova, P., Smith, G. L., and Graham, S. C. (2015) Vaccinia virus protein A49 is an unexpected member of the B-cell Lymphoma (Bcl)-2 protein family. J Biol Chem 290, 5991–6002

10. Morelli, M., Dennis, A. F., and Patton, J. T. (2015) Putative E3 ubiquitin ligase of human rotavirus inhibits NF-kappaB activation by using molecular mimicry to target beta-TrCP. MBio 6

11. Margottin, F., Bour, S. P., Durand, H., Selig, L., Benichou, S., Richard, V., Thomas, D., Strebel, K., and Benarous, R. (1998) A novel human WD protein, h-beta TrCp, that interacts with HIV-1 Vpu connects CD4 to the ER degradation pathway through an F-box motif. Mol Cell 1, 565–574

12. Huang, B., Yang, X. D., Lamb, A., and Chen, L. F. (2010) Posttranslational modifications of NF-kappaB: another layer of regulation for NF-kappaB signaling pathway. Cell Signal 22, 1282–1290

13. Komander, D., and Rape, M. (2012) The ubiquitin code. Annu Rev Biochem 81, 203–229

14. Petroski, M. D., and Deshaies, R. J. (2005) Function and regulation of cullin–RING ubiquitin ligases. Nature reviews. Molecular cell biology 6, 9

15. Kamura, T., Maenaka, K., Kotoshiba, S., Matsumoto, M., Kohda, D., Conaway, R. C., Conaway, J. W., and Nakayama, K. I. (2004) VHL-box and SOCS-box domains determine binding specificity for Cul2-Rbx1 and Cul5-Rbx2 modules of ubiquitin ligases. Genes Dev 18, 3055–3065

16. Yu, Y., Xiao, Z., Ehrlich, E. S., Yu, X., and Yu, X. F. (2004) Selective assembly of HIV-1 Vif-Cul5-ElonginB-ElonginC E3 ubiquitin ligase complex through a novel SOCS box and upstream cysteines. Genes Dev 18, 2867–2872

17. Hilton, D. J., Richardson, R. T., Alexander, W. S., Viney, E. M., Willson, T. A., Sprigg, N. S., Starr, R., Nicholson, S. E., Metcalf, D., and Nicola, N. A. (1998) Twenty proteins containing a C-terminal SOCS box form five structural classes. Proc Natl Acad Sci U S A 95, 114–119

18. Kohroki, J., Nishiyama, T., Nakamura, T., and Masuho, Y. (2005) ASB proteins interact with Cullin5 and Rbx2 to form E3 ubiquitin ligase complexes. FEBS Lett 579, 6796–6802

19. Linossi, E. M., Babon, J. J., Hilton, D. J., and Nicholson, S. E. (2013) Suppression of cytokine signaling: the SOCS perspective. Cytokine Growth Factor Rev 24, 241–248

20. Lewis, R. S., Kolesnik, T. B., Kuang, Z., D’Cruz, A. A., Blewitt, M. E., Masters, S. L., Low, A., Willson, T., Norton, R. S., and Nicholson, S. E. (2011) TLR regulation of SPSB1 controls inducible nitric oxide synthase induction. J Immunol 187, 3798–3805

21. Nishiya, T., Matsumoto, K., Maekawa, S., Kajita, E., Horinouchi, T., Fujimuro, M., Ogasawara, K., Uehara, T., and Miwa, S. (2011) Regulation of inducible nitric-oxide synthase by the SPRY domain- and SOCS box-containing proteins. J Biol Chem 286, 9009–9019

22. Sumner, R. P., Maluquer de Motes, C., Veyer, D. L., and Smith, G. L. (2014) Vaccinia virus inhibits NF-kappaB-dependent gene expression downstream of p65 translocation. J Virol 88, 3092–3102

23. Bhinder, B., and Djaballah, H. (2012) A simple method for analyzing actives in random RNAi screens: introducing the “H Score” for hit nomination & gene prioritization. Comb Chem High Throughput Screen 15, 686–704

24. Rehwinkel, J., Tan, C. P., Goubau, D., Schulz, O., Pichlmair, A., Bier, K., Robb, N., Vreede, F., Barclay, W., Fodor, E., and Reis e Sousa, C. (2010) RIG-I detects viral genomic RNA during negative-strand RNA virus infection. Cell 140, 397–408

25. Unterholzner, L., Sumner, R. P., Baran, M., Ren, H., Mansur, D. S., Bourke, N. M., Randow, F., Smith, G. L., and Bowie, A. G. (2011) Vaccinia virus protein C6 is a virulence factor that binds TBK-1 adaptor proteins and inhibits activation of IRF3 and IRF7. PLoS pathogens 7, e1002247

26. Torres, A. A., Albarnaz, J. D., Bonjardim, C. A., and Smith, G. L. (2016) Multiple Bcl-2 family immunomodulators from vaccinia virus regulate MAPK/AP-1 activation. Journal of General Virology 97, 2346–2351

27. Barrios-Rodiles, M., Brown, K. R., Ozdamar, B., Bose, R., Liu, Z., Donovan, R. S., Shinjo, F., Liu, Y., Dembowy, J., Taylor, I. W., Luga, V., Przulj, N., Robinson, M., Suzuki, H., Hayashizaki, Y., Jurisica, I., and Wrana, J. L. (2005) High-throughput mapping of a dynamic signaling network in mammalian cells. Science 307, 1621–1625

28. Maluquer de Motes, C., Cooray, S., Ren, H., Almeida, G. M., McGourty, K., Bahar, M. W., Stuart, D. I., Grimes, J. M., Graham, S. C., and Smith, G. L. (2011) Inhibition of apoptosis and NF-kappaB activation by vaccinia protein N1 occur via distinct binding surfaces and make different contributions to virulence. PLoS Pathog 7, e1002430

29. Maluquer de Motes, C., Schiffner, T., Sumner, R. P., and Smith, G. L. (2014) Vaccinia virus virulence factor N1 can be ubiquitylated on multiple lysine residues. J Gen Virol 95, 2038–2049

30. Christian, F., Smith, E. L., and Carmody, R. J. (2016) The Regulation of NF-kappaB Subunits by Phosphorylation. Cells 5

31. Sun, Y., and Lopez, C. B. (2017) The innate immune response to RSV: Advances in our understanding of critical viral and host factors. Vaccine 35, 481–488

32. Linossi, E. M., and Nicholson, S. E. (2012) The SOCS box-adapting proteins for ubiquitination and proteasomal degradation. IUBMB Life 64, 316–323

33. Kuang, Z., Lewis, R. S., Curtis, J. M., Zhan, Y., Saunders, B. M., Babon, J. J., Kolesnik, T. B., Low, A., Masters, S. L., Willson, T. A., Kedzierski, L., Yao, S., Handman, E., Norton, R. S., and Nicholson, S. E. (2010) The SPRY domain-containing SOCS box protein SPSB2 targets iNOS for proteasomal degradation. J Cell Biol 190, 129–141

34. Feng, Y., Pan, T. C., Pant, D. K., Chakrabarti, K. R., Alvarez, J. V., Ruth, J. R., and Chodosh, L. A. (2014) SPSB1 promotes breast cancer recurrence by potentiating c-MET signaling. Cancer Discov 4, 790–803

35. Wang, D., Li, Z., Messing, E. M., and Wu, G. (2005) The SPRY domain-containing SOCS box protein 1 (SSB-1) interacts with MET and enhances the hepatocyte growth factor-induced Erk-Elk-1-serum response element pathway. J Biol Chem 280, 16393–16401

36. Filippakopoulos, P., Low, A., Sharpe, T. D., Uppenberg, J., Yao, S., Kuang, Z., Savitsky, P., Lewis, R. S., Nicholson, S. E., Norton, R. S., and Bullock, A. N. (2010) Structural basis for Par-4 recognition by the SPRY domain- and SOCS box-containing proteins SPSB1, SPSB2, and SPSB4. J Mol Biol 401, 389–402

37. Masters, S. L., Yao, S., Willson, T. A., Zhang, J. G., Palmer, K. R., Smith, B. J., Babon, J. J., Nicola, N. A., Norton, R. S., and Nicholson, S. E. (2006) The SPRY domain of SSB-2 adopts a novel fold that presents conserved Par-4-binding residues. Nat Struct Mol Biol 13, 77–84

38. Liu, S., Iaria, J., Simpson, R. J., and Zhu, H. J. (2018) Ras enhances TGF-beta signaling by decreasing cellular protein levels of its type II receptor negative regulator SPSB1. Cell Commun Signal 16, 10

39. Liu, S., Nheu, T., Luwor, R., Nicholson, S. E., and Zhu, H. J. (2015) SPSB1, a Novel Negative Regulator of the Transforming Growth Factor-beta Signaling Pathway Targeting the Type II Receptor. J Biol Chem 290, 17894–17908

40. Wang, F., Fu, X., Chen, P., Wu, P., Fan, X., Li, N., Zhu, H., Jia, T. T., Ji, H., Wang, Z., Wong, C. C., Hu, R., and Hui, J. (2017) SPSB1-mediated HnRNP A1 ubiquitylation regulates alternative splicing and cell migration in EGF signaling. Cell Res 27, 540–558

41. Perkins, N. D. (2006) Post-translational modifications regulating the activity and function of the nuclear factor kappa B pathway. Oncogene 25, 6717–6730

42. Pautz, A., Art, J., Hahn, S., Nowag, S., Voss, C., and Kleinert, H. (2010) Regulation of the expression of inducible nitric oxide synthase. Nitric Oxide 23, 75–93

43. Croker, B. A., Krebs, D. L., Zhang, J. G., Wormald, S., Willson, T. A., Stanley, E. G., Robb, L., Greenhalgh, C. J., Forster, I., Clausen, B. E., Nicola, N. A., Metcalf, D., Hilton, D. J., Roberts, A. W., and Alexander, W. S. (2003) SOCS3 negatively regulates IL-6 signaling in vivo. Nat Immunol 4, 540–545

44. Kershaw, N. J., Murphy, J. M., Liau, N. P., Varghese, L. N., Laktyushin, A., Whitlock, E. L., Lucet, I. S., Nicola, N. A., and Babon, J. J. (2013) SOCS3 binds specific receptor-JAK complexes to control cytokine signaling by direct kinase inhibition. Nat Struct Mol Biol 20, 469–476

45. Frobose, H., Ronn, S. G., Heding, P. E., Mendoza, H., Cohen, P., Mandrup-Poulsen, T., and Billestrup, N. (2006) Suppressor of cytokine Signaling-3 inhibits interleukin-1 signaling by targeting the TRAF-6/TAK1 complex. Mol Endocrinol 20, 1587–1596

46. Seki, Y., Hayashi, K., Matsumoto, A., Seki, N., Tsukada, J., Ransom, J., Naka, T., Kishimoto, T., Yoshimura, A., and Kubo, M. (2002) Expression of the suppressor of cytokine signaling-5 (SOCS5) negatively regulates IL-4-dependent STAT6 activation and Th2 differentiation. Proc Natl Acad Sci U S A 99, 13003–13008

47. Kario, E., Marmor, M. D., Adamsky, K., Citri, A., Amit, I., Amariglio, N., Rechavi, G., and Yarden, Y. (2005) Suppressors of cytokine signaling 4 and 5 regulate epidermal growth factor receptor signaling. J Biol Chem 280, 7038–7048

48. Nicholson, S. E., Metcalf, D., Sprigg, N. S., Columbus, R., Walker, F., Silva, A., Cary, D., Willson, T. A., Zhang, J. G., Hilton, D. J., Alexander, W. S., and Nicola, N. A. (2005) Suppressor of cytokine signaling (SOCS)-5 is a potential negative regulator of epidermal growth factor signaling. Proc Natl Acad Sci U S A 102, 2328–2333

49. Kedzierski, L., Tate, M. D., Hsu, A. C., Kolesnik, T. B., Linossi, E. M., Dagley, L., Dong, Z., Freeman, S., Infusini, G., Starkey, M. R., Bird, N. L., Chatfield, S. M., Babon, J. J., Huntington, N., Belz, G., Webb, A., Wark, P. A., Nicola, N. A., Xu, J., Kedzierska, K., Hansbro, P. M., and Nicholson, S. E. (2017) Suppressor of cytokine signaling (SOCS)5 ameliorates influenza infection via inhibition of EGFR signaling. Elife 6

50. Kamizono, S., Hanada, T., Yasukawa, H., Minoguchi, S., Kato, R., Minoguchi, M., Hattori, K., Hatakeyama, S., Yada, M., Morita, S., Kitamura, T., Kato, H., Nakayama, K., and Yoshimura, A. (2001) The SOCS box of SOCS-1 accelerates ubiquitin-dependent proteolysis of TEL-JAK2. J Biol Chem 276, 12530–12538

51. Strebovsky, J., Walker, P., Lang, R., and Dalpke, A. H. (2011) Suppressor of cytokine signaling 1 (SOCS1) limits NFkappaB signaling by decreasing p65 stability within the cell nucleus. FASEB J 25, 863–874

52. Barretina, J., Caponigro, G., Stransky, N., Venkatesan, K., Margolin, A. A., Kim, S., Wilson, C. J., Lehar, J., Kryukov, G. V., Sonkin, D., Reddy, A., Liu, M., Murray, L., Berger, M. F., Monahan, J. E., Morais, P., Meltzer, J., Korejwa, A., Jane-Valbuena, J., Mapa, F. A., Thibault, J., Bric-Furlong, E., Raman, P., Shipway, A., Engels, I. H., Cheng, J., Yu, G. K., Yu, J., Aspesi, P., Jr., de Silva, M., Jagtap, K., Jones, M. D., Wang, L., Hatton, C., Palescandolo, E., Gupta, S., Mahan, S., Sougnez, C., Onofrio, R. C., Liefeld, T., MacConaill, L., Winckler, W., Reich, M., Li, N., Mesirov, J. P., Gabriel, S. B., Getz, G., Ardlie, K., Chan, V., Myer, V. E., Weber, B. L., Porter, J., Warmuth, M., Finan, P., Harris, J. L., Meyerson, M., Golub, T. R., Morrissey, M. P., Sellers, W. R., Schlegel, R., and Garraway, L. A. (2012) The Cancer Cell Line Encyclopedia enables predictive modelling of anticancer drug sensitivity. Nature 483, 603–607

53. Schuliga, M. (2015) NF-kappaB Signaling in Chronic Inflammatory Airway Disease. Biomolecules 5, 1266–1283

54. Hayashi, R., Wada, H., Ito, K., and Adcock, I. M. (2004) Effects of glucocorticoids on gene transcription. Eur J Pharmacol 500, 51–62

55. Edwards, M. R., Bartlett, N. W., Clarke, D., Birrell, M., Belvisi, M., and Johnston, S. L. (2009) Targeting the NF-kappaB pathway in asthma and chronic obstructive pulmonary disease. Pharmacol Ther 121, 1–13

56. Schaller, T., Ocwieja, K. E., Rasaiyaah, J., Price, A. J., Brady, T. L., Roth, S. L., Hue, S., Fletcher, A. J., Lee, K., KewalRamani, V. N., Noursadeghi, M., Jenner, R. G., James, L. C., Bushman, F. D., and Towers, G. J. (2011) HIV-1 capsid-cyclophilin interactions determine nuclear import pathway, integration targeting and replication efficiency. PLoS Pathog 7, e1002439

57. Zufferey, R., Nagy, D., Mandel, R. J., Naldini, L., and Trono, D. (1997) Multiply attenuated lentiviral vector achieves efficient gene delivery in vivo. Nat Biotechnol 15, 871–875

58. Odon, V., Georgana, I., Holley, J., Morata, J., and Maluquer de Motes, C. (2018) A novel class of viral Ankyrin proteins targeting the host E3 ubiquitin ligase Cullin-2. J Virol

59. Gao, D., Wu, J., Wu, Y. T., Du, F., Aroh, C., Yan, N., Sun, L., and Chen, Z. J. (2013) Cyclic GMP-AMP synthase is an innate immune sensor of HIV and other retroviruses. Science 341, 903–906

60. Georgana, I., Sumner, R. P., Towers, G. J., and Maluquer de Motes, C. (2018) Virulent poxviruses inhibit DNA sensing by preventing STING activation. J Virol 1. 124.

61. Chen, R. A., Ryzhakov, G., Cooray, S., Randow, F., and Smith, G. L. (2008) Inhibition of IkappaB kinase by vaccinia virus virulence factor B14. PLoS pathogens 4, e22

62. Maluquer de Motes, C., and Smith, G. L. (2017) Vaccinia virus protein A49 activates Wnt signalling by targetting the E3 ligase beta-TrCP. J Gen Virol

63. Thurston, T. L., Ryzhakov, G., Bloor, S., von Muhlinen, N., and Randow, F. (2009) The TBK1 adaptor and autophagy receptor NDP52 restricts the proliferation of ubiquitin-coated bacteria. Nat Immunol 10, 1215–1221

